# Motility mediates satellite formation in confined biofilms

**DOI:** 10.1101/2022.12.16.520725

**Authors:** Mireia Cordero, Namiko Mitarai, Liselotte Jauffred

**Affiliations:** The Niels Bohr Institute, University of Copenhagen, Blegdamsvej 17, DK-2100 Copenhagen O, Denmark

## Abstract

Bacteria have spectacular survival capabilities and can spread in many, vastly different environments. For instance, when pathogenic bacteria infect a host, the cells expand through proliferation and squeezing through narrow pores and elastic matrices. However, the exact role of surface structures and matrix elasticity in colony expansion and morphogenesis is still largely unknown. Here we show how *satellite* colonies emerge around biofilms embedded in semi-soft agar in controlled *in vitro* assays. We tested how extra-cellular structures – important for biofilm formation and motility – control this morphology. Moreover, we identify the range of extra-cellular matrix elasticity, where this morphology is possible. When paralleled with mathematical modelling, our results demonstrate that satellite formation allows bacterial communities to spread faster. We anticipate that this strategy is important to speed up expansion in various environments while retaining the close interactions and protection provided by the community.

## Introduction

While many bacteria species can grow as free-swimming cells in planktonic mode, they will often adhere to a substrate or small enclosure to form dense communities, i.e., biofilms^1^. The biofilm community offers protection from local threats, e.g., against the shear flow by attaching to a solid substrate or by protecting the inner cells from the immune system, toxins, or bacteriophages^2,3^. Also, the local, high cell density helps bacteria share necessary chemicals within the community^4^. The importance of biofilm formation is also reflected in the fact that bacteria have many genes contributing to cell-adhesive structures, such as exo-polysaccharides (EPS), fimbriae, and flagella (see reference^5^ for a review). These biofilms often grow – as the name indicates – as quasi-two-dimensional (2D) colonies on substrates, but also as three-dimensional (3D) communities in habitats, such as gels, tissues (e.g., human guts), and soils. Despite this high prevalence, we still lack a full understanding of how dynamic morphologies of 3D biofilms are controlled.

When a biofilm grows as a dense colony, the surface expands outwards as cells are proliferating, with a doubling time close to what we know from well-mixed liquid cultures. Behind the fast-growing pioneering cells on the front, is a quiescent region, where proliferation is slowed down dramatically due to, e.g., space constraints^6^ and metabolite limitations^7,8^. The resulting expansion of the colony surface makes the population grow linearly over time^8,9^, in contrast to the exponential growth in liquid cultures. Therefore, the motility of cells plays a crucial role to speed up colony expansion. It is well-established by swimming assays in soft agar that chemotaxis is enhanced by nutrient shortage^10–12^ or attractant gradients^13^. Chemotaxis accelerates expansion dynamics of populations by allowing access to more nutrient^14^. In liquid media, the chemotaxis of the model bacterium *Escherichia coli* is driven by swimming, i.e., bundling and propelling of flagella, which gives rise to a run-and-tumble motion. In visco-elastic media, bacterial migration is restricted more and more as elasticity is raised and pore size diminished^15^ and the chemotaxis has been reported to be reversely proportional to the elasticity of the medium^12^. Therefore, above a certain threshold of medium elasticity, biofilm expansion is solely growth-driven. Furthermore, in very elastic matrices, the stress at the interface (between colony and media) gives rise to the internal ordering of cells^16^. But what happens in the intermediate regions, where the agar is semi-soft? Is it possible that cells can both retain biofilm characteristics but still speed up the expansion to enhance colonization?

Here we use mono-clonal *E. coli* colonies embedded in a semi-solid agarose matrix^2,17^ as simple models of 3D biofilm formation. We use a combination of experiments and mathematical modelling to show how 3D bacterial biofilms can join growth and flagella-based motility to colonize their local environment. We investigate how the spreading depends on the expression of extra-cellular structures and matrix elasticity. We demonstrate that indeed 3D biofilm expansion – driven by the combination of growth and motility – gives rise to satellite colony formation and acceleration of the population expansion to be super-linear over time.

## Results

### Satellite colonies form around 3D biofilms

To mimic 3D biofilm evolvement in natural settings, *E. coli* MG1566 wild-type (wt) single cells were embedded in low concentration (≈10 cells pr. ml) in a soft agarose (0.3% agarose) matrix and incubated for 15 hours at 37°C in a minimal defined medium composed of M63 supplemented with 20 *μ*m/ml glucose (M63+glu). This protocol (see Materials and methods for details) resulted in confined, mono-clonal 3D biofilms on the order of a few hundred *μ*m in diameter. In parallel, we performed a similar experiment using a rich Luria-Bertani (LB) medium and a shorter incubation time of 13 hours to account for differences in doubling time in the two media: 27.8 min (LB) vs. 68.4 min (M63+glu), see Supplementary figure S1.

The resulting biofilms in figure 1A grew as compact quasi-spherical colonies with a mountainous surface of multiple protrusions, as in the first column of images in figure 1A denoted as wt −. However, for the majority of 3D biofilms, 34/40 (M63+glu) and 17/18 (LB), these protrusions seem to have escaped the colony as small stationary satellite colonies, as shown in figure 1A and denoted as wt+. Both morphology types occurred for biofilms grown in both the minimal (M63+glu) and the rich media (LB). It is worth noticing that these two morphologies are not determined by any obvious variations between experiments, as we found wt− and wt+ side by side in the same culture wells.

**Figure 1.**
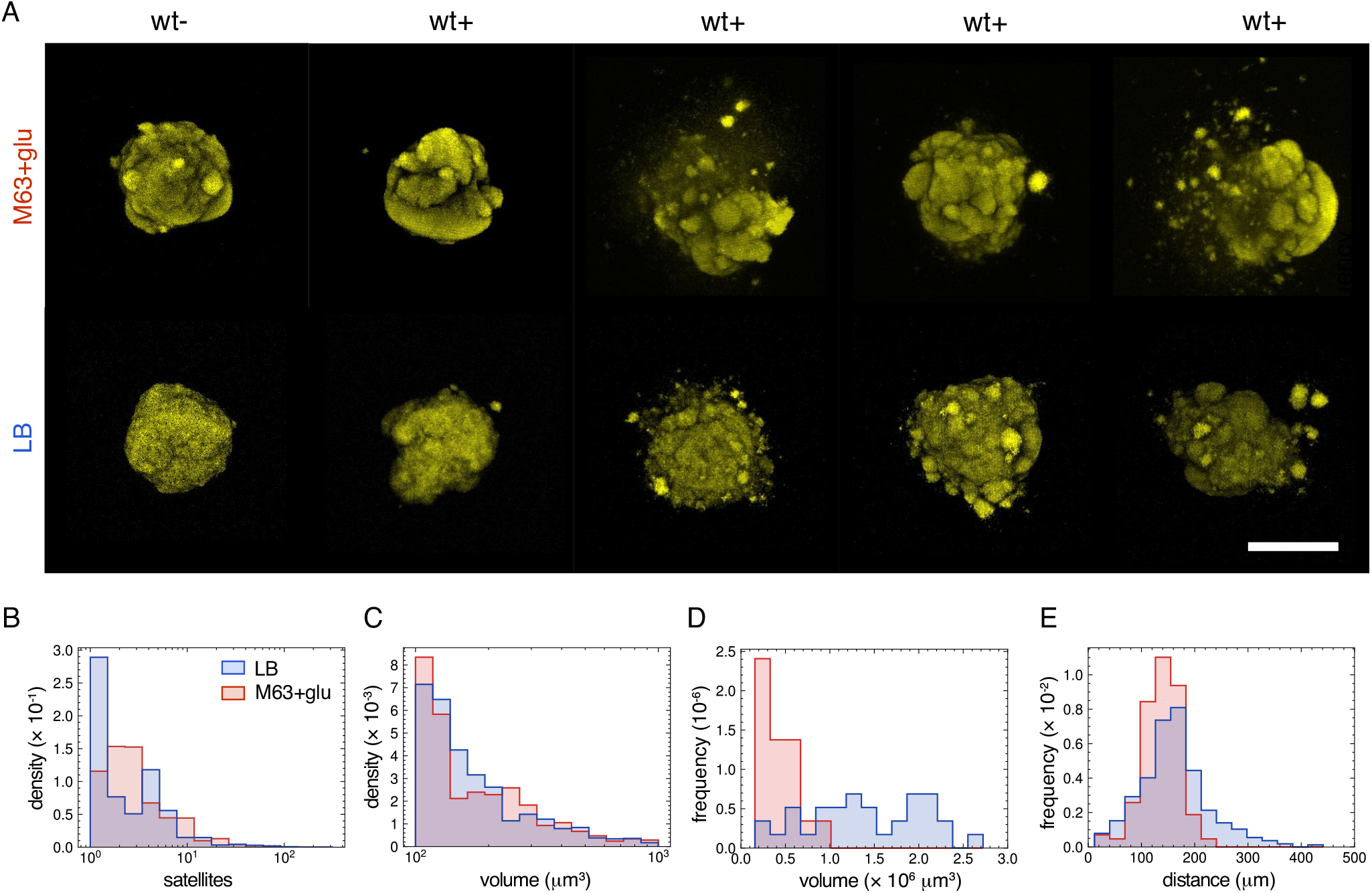
**A:** Satellite morphology in mature 3D biofilms. Examples of pseudo-colored 3D biofilms (maximum intensity projections) in 0.3% agarose and minimal (M63+glu) or rich medium (LB). These examples shows inhomogeneity of morphologies without satellites (wt −) or with satellites (wt+). The scale bar corresponds to 200 *μ*m. **B:** Distributions of the number of satellites pr. biofilm (log-scale) in either LB (N=17) and M63+glu (N=34). **C:** Distributions of satellite volumes (log-scale) in either LB (N=149) or in M63+glu (N=574). **D:** Distribution of volumes of the main colonies in either LB (N=17) and M63+glu (N=34). **E:** Distributions of distances from the center-of-mass of satellites to the center-of-mass of the main colony in either LB (N=17) and M63+glu (N=34).

From the subset of 3D biofilms with satellite morphology (wt+), we measured the distributions of number of satellites, their volumes and distances to the main colony, as well as the main colony volumes, for both minimal (blue) and rich (red) medium (Figure 1B-E). The distribution of the number of satellites per 3D biofilm (Figure 1B) shows that the majority have less than 10 satellites. Furthermore, we find that satellites with smaller volumes have higher prevalence as seen from the distribution of satellite volumes on a semi-logarithmic plot in Figure 1C. The typical satellite size (*<* 10^3^ *μ*m^3^) is about 3 orders of magnitude smaller than the average volumes of the main colonies, which are (1.3 *±* 0.6) ·10^6^ *μ*m^3^ (M63+glu) and (4 *±* 1) · 10^5^ *μ*m^3^ (LB). The distribution of the main colony size is given in figure 1D. Finally, Figure 1E shows the distributions of distances between center-of-masses of the satellites and their respective main colony. The average distances are 161 *±* 64 *μ*m and 137 *±* 35 *μ*m for minimal (M63+glu) and rich (LB) medium, respectively.

A possible mechanism behind satellite formation is the combination of bacterial cell-cell interaction, motility, and the growth of the cells: A few cells can move away from the surface of the main colony escaping from their attachment to migrate to a different position in the gel, and each of them starts to grow into a new satellite colony. Therefore, in the following, we investigate how bacterial surface structures, as well as the elasticity of the substrate, affect the emergence of satellite colonies. The first is achieved by studying mutants with deletions of structures important for adhesion and motility, and the latter by modifying agarose concentration in the extracellular matrix.

### Deletion of extracellular structures causes loss of satellite morphology

One of the factors governing the morphology of 3D biofilms and the appearance of satellites must be the complex interplay between cell envelope components and the extracellular matrix. So, bacterial cell-adhesive structures, such as exo-polysaccharides (EPS), fimbriae, and flagella are obvious components that can modulate the biofilm morphology. Therefore, we investigated a collection of mutants lacking diverse bacterial surface structures. Information on all strains used in this study is found in Table 1. Figure 2A sketches the parental strain (wt) and the suppressed structures: flagella, type I pili, colanic acid, curli fimbriae and antigen 43. To examine the influence of these surface components, we examined the 3D biofilm morphology after knock-out deletion and compared it with the wt colony morphology in Figure 2B.

**Table 1.** *E. coli* strains used in the study

| Name | Strain | Relevant characteristics | Reference |
| --- | --- | --- | --- |
| wt | MS613 | MG1655 K-12 reference (F-lambda- ilvG- rfb-50 rph-1) strain | 57 |
| Δflu | MS427 | MG1655 Δflu | 63 |
| Δfim | MS428 | MG1655 Δfim | 22 |
| ΔfluΔfim | MS528 | MG1655 Δflu,Δfim | 22 |
| Δcps | RMV340 | MG1655cps::tet | 64 |
| ΔcsgAB | RMV612 | MG1655csgAB::kan | 58 |
| ΔfliC | NM109 | MG1655fliC::kan | This study |

**Figure 2.**
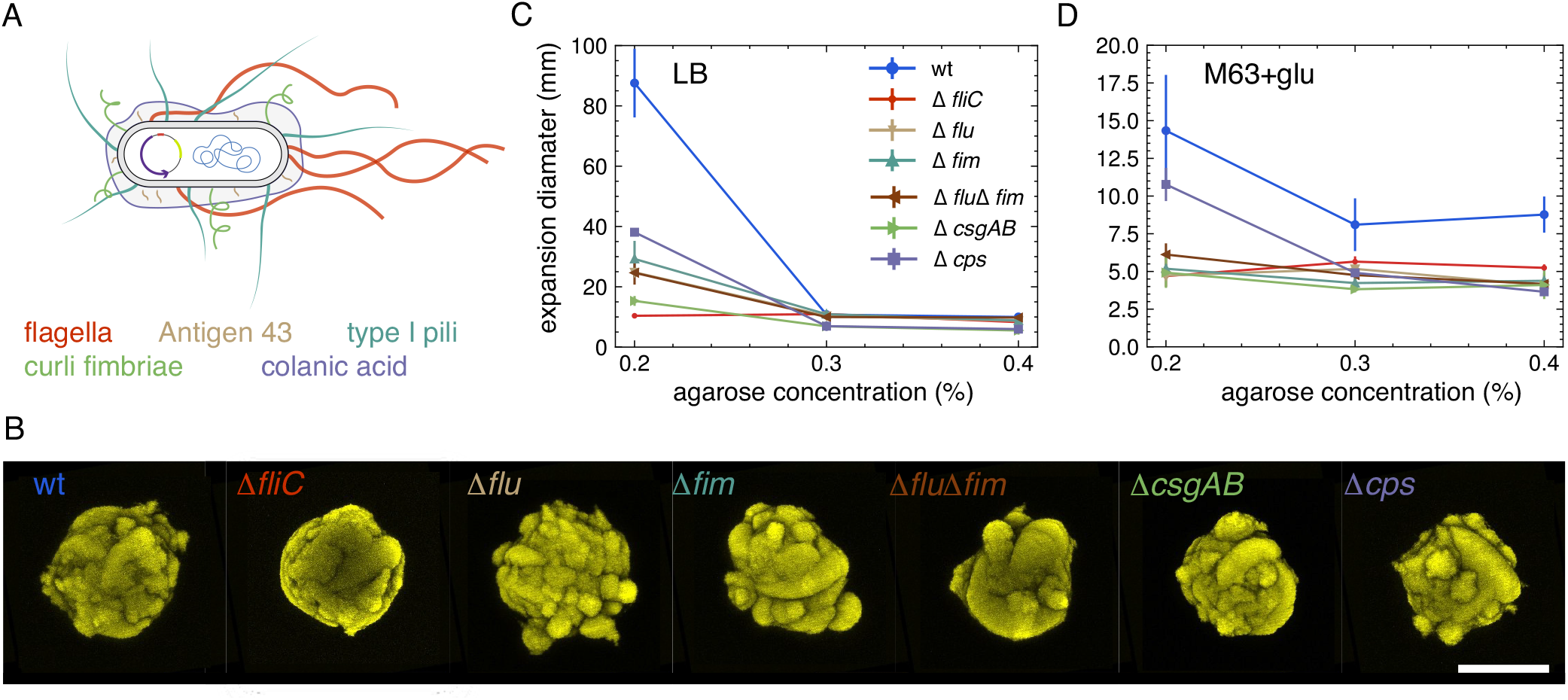
**A:** Sketch of parental (wt) *E.coli* strain (not to scale). The sketched elements includes the genome, the fluorescence-carrying plasmid and the extracellular structures: flagella (red), antigen 43 (beige), type I pili (dark green), curli fimbriae (light green), colanic acid (purple). **B:** Morphology of mutant 3D biofilms. Examples of pseudo-colored fluorescent biofilms (maximum intensity projections) grown in M63+glu with 0.3% agarose. Scale bar corresponds to 200 *μ*m and color coding is the same as in (A). **C+D:** Swimming assay in soft agar. Diameters of expansion zones after 24 hours of incubation vs. agarose concentrations in LB (C) and M63+glu (D).

Flagella are helical filaments responsible for some of the locomotion mechanisms adopted by bacteria. By expressing flagella, cells can self-propel in liquids (swimming) and on surfaces (swarming)^18^. Furthermore, flagella have been shown to play a role in cell–substratum (e.g. initial biofilm formation^19^) and cell-cell interactions^5^. However, the role of these structures in bacterial motility in confined environments remains laregely unknown^20^. We tested a Δ*fliC* mutant and did not find any satellite formation (N=21). We also assessed a Δ*fim* mutant without type I pili, i.e., a hair-like structures crucial for cell-cell adhesion^19^. Surprisingly, the morphology of these mutants is also indistinguishable from that of wt − (N=21).

The Δ*flu* mutant has a deletion in the gene encoding the auto-transporter protein antigen 43, which comes in high copy numbers (up to 50,000 pr. cell^21^). Antigen 43 favors cell-cell interactions, e.g., chain formation, and is referred to as a *handshake* protein^22–24^. Given the inverse regulation of pili and antigen 43^25^, we tested both Δ*flu* (N=21) and the double mutant Δ*flu*Δ*fim*(N=22) and found that both of them display a loss of satellite morphology.

We also tested the EPS colonic acid mutant, Δ*cps* (N=17), as well as the Δ*csgAB* (N=18) with a deletion in the gene encoding a fibrous surface protein, curli fimbriae, important for both cell-cell and cell-substrate adhesion^26–30^. Even though *cps* and *csgAB* predominantly are expressed at ambient temperature^28,31^, we found wt+ morphology to be lost in both cases. In summary, we found that the satellite morphology (wt+) is lost in the confined biofilms of all tested mutants.

### Deletion of extracellular structures reduces motility

Except for the flagella deletion that disables the most common mechanisms of bacterial motility (swimming/swarming), the effect of the other tested surface structures on bacterial confined motility is less obvious. To investigate this further, we performed classical motility assays in soft agar for all strains used in this study. The assay was done using both rich (LB) and minimal media (M63+glu) with varying agarose concentrations (0.2%-0.4%).

Figure 2C shows that the wt is the most motile strain in the soft (0.2% agarose) rich agar (LB), with an average distance reached of about 3*×* the one of the second fastest, Δ*cps*. All strains are found to be motile except the non-flagellated mutant, Δ*fliC*, but with large variations in speed. This is in accordance with the fact that many of the bacterial surface structures display interdependent regulation^32^. At higher substrate stiffness (0.3%-0.4% agarose), where swimming is no longer the main driver of motility, the average moving distances are considerably reduced for all strains.

We also tested all strains’ behavior in a minimal medium (M63+glu), as shown in Figure 2D. In the soft minimal agar (0.2% agarose) the motility of all strains is reduced, as can be seen by comparing the maximum distance reached by wt: (14.3 *±* 3.7) cm vs. (87.5 *±*11.4) cm in rich medium (LB). The reduced motility in minimal media is consistent with the previous studies: It has been attributed to the considerable metabolic cost of producing flagella^33,34^, while a recent study^13^ has shown that the chemotaxis is strongly reduced when there is no supplement attractant, even if the primary carbon source is also an attractant. Again, at greater agarose concentrations (0.3%-0.4% agarose) motility reduces to a non-detectable level for all other than the wt.

Overall, the soft agar assay suggests that all the mutant strains tested here have reduced motility compared to the wt strain. This is in line with the view that the satellite colonies form as a compromise between their inherent motility and the mechanical constraints imposed by the environment.

### Elasticity of extra-cellular matrix controls satellite emergence

Another indisputable factor that affects both bacterial motility and biofilm morphology is the elasticity of the matrix that encloses the cells. To explore the colony morphology dependence on this environmental factor, we limited our analysis to a comparison between the highly motile wt and the non-flagellated mutant (Δ*fliC*) that was non-motile in all tested experimental settings. We investigated mono-clonal colonies of both strains in varying agarose concentration (0.25%-0.35%) in rich (LB) and minimal (M63+glu) medium.

As already mentioned, we found satellite morphology (wt+) among the 3D biofilms in 0.3% agarose with frequencies of 34/40 and 17/18 for the wt strain in minimal and rich medium, respectively. Namely, the satellite morphology is conserved despite changes in nutrient composition and possible slight changes in elasticity between the experimental samples^35^. Examples of the resulting 3D biofilms are given in figure 3 for both minimal medium (M63+glu) and rich medium (LB).

**Figure 3.**
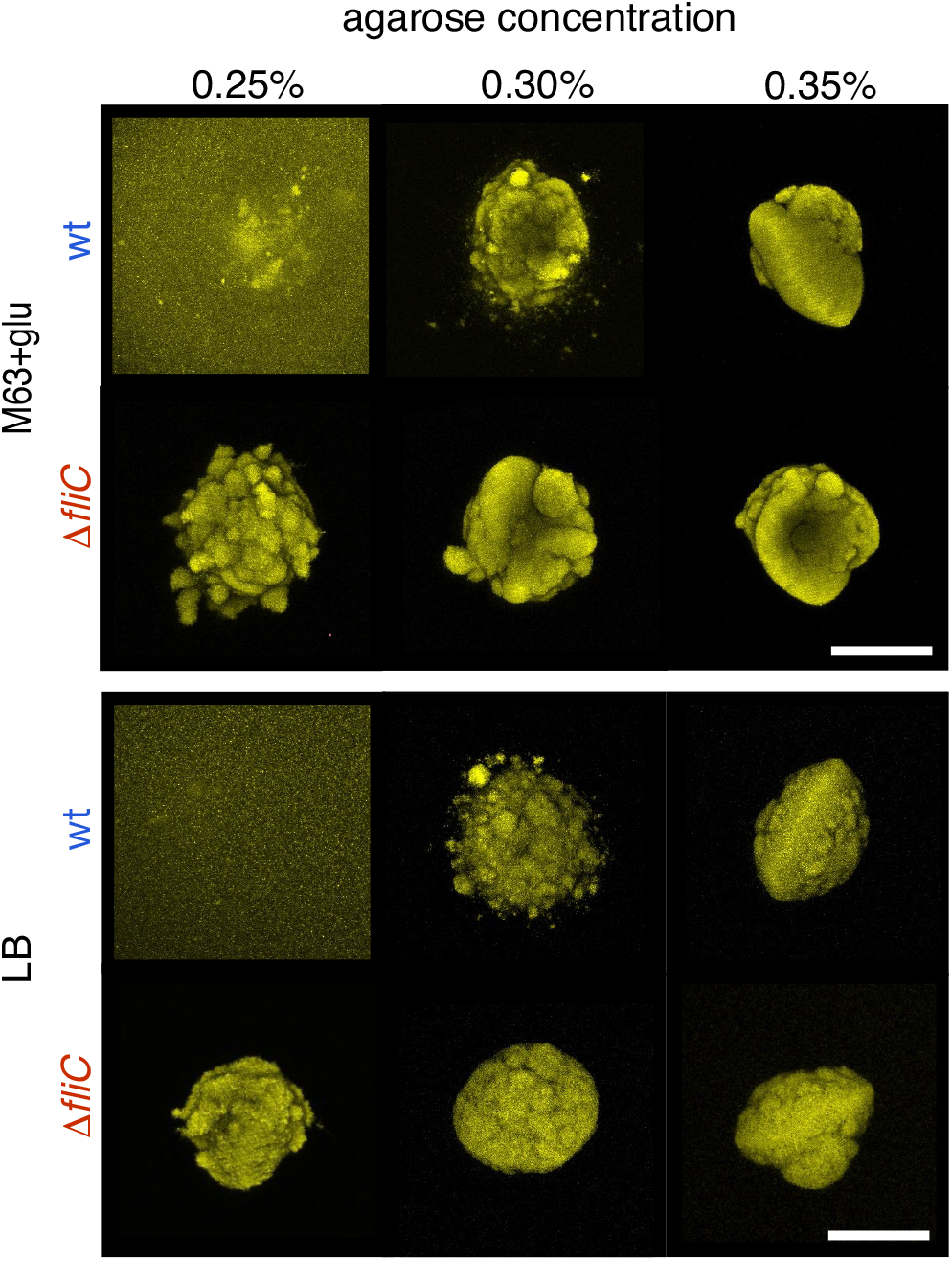
Effect on matrix elasticity on 3D biofilm morphology. Examples of pseudo-colored fluorescent parental (wt) or flagella mutant (Δ*fliC*) biofilms (maximum intensity projections) grown in either minimal (M63+glu) or rich (LB) medium at various agarose concentrations. The scale bar corresponds to 200 *μ*m.

When reducing the agarose concentration (0.25%) the distinct satellite morphology is lost for wt, and general spreading of the cells inside the gel is observed (Figure 3, 0.25%). On the other hand, when stiffness was increased (0.35%), 3D biofilms grew under greater confinement, resulting in the loss of wt+ morphology. In contrast, the non-motile strain (Δ*fliC*) always grows as single compact colonies at all tested agarose concentrations. We still observe the agarose concentration dependence on the morphology, as at 0.25% agarose concentration the surface of Δ*fliC* biofilms appears more loosely connected than at higher agarose concentrations.

Noticeably, colony surfaces become smoother as matrix stiffness increases for both strains (Figure 3, 0.35%). In some cases (7/18 in M63-glu and 6/9 in LB), we even found slightly oblate colony shapes. This shape change depends on the elasticity of the environment, such that for higher stiffness (*>* 0.4% agarose) all tested 3D biofilms grow as single oblate colonies. This is in agreement with earlier reports^36^ and examples are shown in Supplementary figure S2. Recently, this shape morphology has been suggested to originate from the stiffness contrast between the biofilms and the environment^16^.

These findings all together suggest that the emergence of satellites is a transitional state between the morphologies: i) where cells can swim – more or less freely – through the extra-cellular matrix, and ii) where cells are strictly confined.

### Satellite formation possibly speeds up colonization

We explored this question through a *simple* mathematical model that reproduced the 3D morphology of the confined bacterial biofilms. For this purpose, we modified the well-known Eden growth model, which is a lattice model, where the surface cells can divide to occupy empty next nearest neighbor sites^37,38^. Our modification allows for occasional escapes of single cells from the main colony. In other words, surface cells do not only grow but can also migrate to another empty site in the lattice. We do not simulate the cells’ trajectory. Instead, we implemented the migration as a jump to the final position, which is chosen randomly by a radial Gaussian probability distribution centered around the starting site. This simplification is based on the assumption that the durations of migrations on lengths small than 100 *μ*m is much faster than the doubling time of wt. Therefore, the model is essentially controlled by two parameters. The first is the rate of jumps, *k*_*s*_, when the cell doubling rate, *k*, is set to unity. The second is the standard deviation, *σ*, of the Gaussian distribution of possible new sites to jump to. We ran the simulation until a million jump/division events. Thus, the resulting *in silico* biofilms have widths of about 100 *μ*m assuming i) colonies are compact and ii) one occupied lattice site corresponds to one cell volume (∼ 1 *μ*m^3^).

Supplementary figure S3 depicts examples of 3D biofilms with *σ* ∈{ 2, 5, 7, 10} and *k*_*s*_ ∈ {0.001, 0.05, 0.1, 0.2}. At low *σ* and *k*_*s*_, the biofilms are compact with smooth surfaces and as the distances (i.e. *σ*) rise, small protrusions appear on the surface followed by satellite breakouts. As anticipated, the number of satellites also increases as the jump frequency (i.e. *k*_*s*_) increases and ultimately (*σ* = 10 and *k*_*s*_ = 0.2) the simulated system is no longer confined. Instead, the cells spread mimicking the experimental results of wt in the soft agar assay (figure 3, 0.25%).

Figure 4A shows selected runs from Supplementary figure S3, where the smooth and compact biofilm (red) mimics the non-motile Δ*fliC* biofilm (e.g. figure 2B) and the three others (blue) different wt biofilms (e.g. figure 1A). As a sanity check, we analysed our *in silico* 3D biofilms (N=30 of each) in the exact same way as in figure 1B-E and found similar trends, as found in Supplementary figure S4. With this we explored time evolution, figure 4B, and found that the biofilms with wt+ morphology (*σ* = 5 and *k*_*s*_ = 0.01, *σ* = 7 and *k*_*s*_ = 0.01, and *σ* = 7 and *k*_*s*_ = 0.01) clearly grew faster than the compact biofilms (*σ* = 2 and *k*_*s*_ = 0.01). Hence, these results predict a view, where satellite outbreaks speed up the colonization of the extra-cellular matrix.

**Figure 4.**
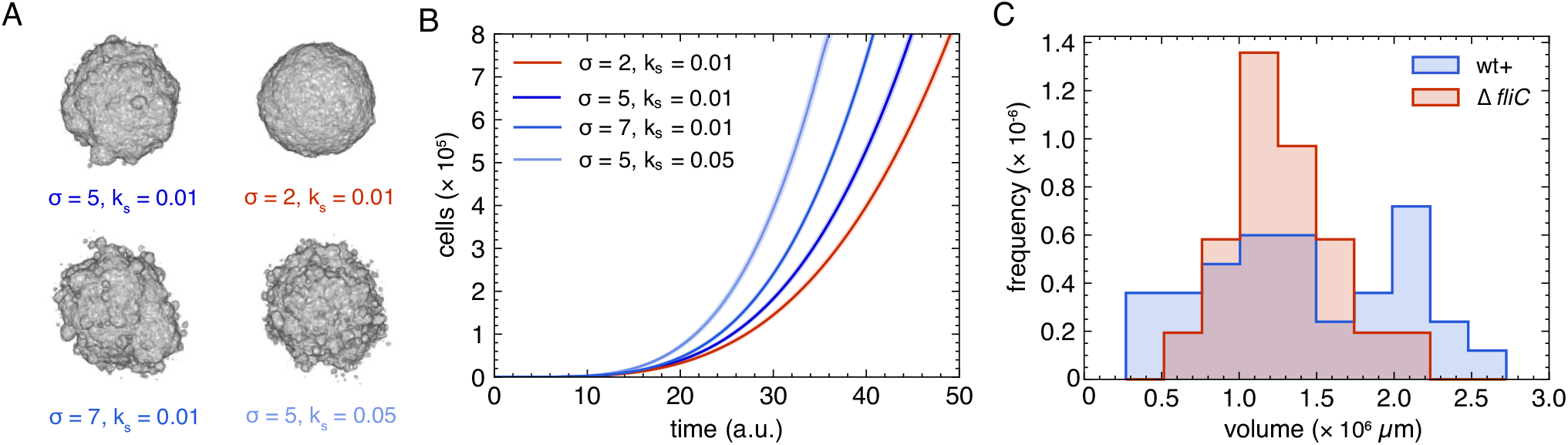
Population size and time evolvement. **A:** Effect of frequency, *k*_*s*_, and distance, *σ*, of jumps in the simulations of 3D biofilms after 10^6^ division/jump events; corresponding to motile (blue) and non-motile (red) biofilms. **B:** Time-evolution of each of the four examples (N=25) given in (A), where time is defined as detailed in Materials and methods. The shaded region corresponds to *±* SD. **C:** Distribution of total biofilm volumes in minimal (M63+glu) medium of wt+ (N=34) and the flagella mutant Δ*fliC* (N=21). The wt+ corresponds to the sum of volumes in figure 1C+D (M63+glu).

To test this prediction, we compared the experimental distributions of final total volumes of wt+ (blue) and Δ*fliC* (red) in figure 4C. We find that the occurrence of satellites results in a more widespread distribution of volumes compared to the non-motile strain (Δ*fliC*). Surprisingly, there are also wt+ biofilms with smaller volumes even though their doubling time may slightly faster than that of Δ*fliC*: 68.4 min vs. 72.9 min (Supplementary figure S1). While the wide distribution makes it hard to compare the average values in the current sample size, the wider distribution with more satellite colonies is in qualitative accordance with the distributions of the simulated 3D biofilms (Supplementary figure S5).

## Discussion

Even though the flagella-mediated motility of *E. coli* is well-described by the run-and-tumble model (see^39^ for a review), we still lack the full understanding of how bacteria migrate through porous or visco-elastic media. Several theoretical works concerning flagella-based motility describe how the migration can be slowed down when colliding with structures of various sizes^40–42^. Moreover, in a recent study, the individual trajectories of bacteria in a porous medium were imaged and the velocity was found to be highly dependent on pore size^15^. Altogether, these reports indicate that *E. coli* moves through a porous medium constantly alternating between stalling (i.e. being trapped), re-orientation, and swimming. In accordance with this, we find that flagella are necessary for migration through the various semi-soft media used in this study (e.g. figure 3). However, the behaviour we unravel is still very different from swimming in soft agar (figure 2C+D) as the main colonies (as well as the satellites) in 0.3% agarose have sharply defined surfaces, indicating that cell outbreak is a rare event. Also, we did not see any signs of the collective colony detachments, recently reported in mature biofilms^43–45^.

Given these observations, we propose that the satellite colonies are formed by the rare – but not too rare – detachment and migration of a single surface cell, followed by stalling, and sub-sequent growth. So, the volumes of satellite colonies indicate when, in the history of the biofilm, the excursion occurred. Furthermore, the size also tells us that the dispersal of the founder cell was long enough to allow the sub-cluster to form before it is annexed to the main colony. We observe that larger satellites (*>* 500 *μ*m^3^) are much less common (figure 3 wt LB 0.3% for an example) than smaller ones (figure 1B), which is consistent with the proposed mechanism.

Our lattice model, which is based on an Eden growth model with rare jumping events added, successfully produced 3D biofilms with satellite morphologies in a specific parameter range. The model predicted that the distribution of total colony volumes would be more widespread when the satellite formation is more frequent (e.g. compare *σ* = 5 and *k*_*s*_ = 0.05 vs. *σ* = 2 and *k*_*s*_ = 0.01 in figure S5). This is consistent with our experimental findings, where we find the distribution of total volumes of the flagella mutant (Δ*fliC*) to be much more narrow than for the parental (wt) strain (figure 4C).

Hallatscheck and co-workers^46,47^ have theoretically demonstrated the impact of long-range dispersal in population expansion, especially when the dispersal distance distribution has a somewhat fat tail. In our model, the distribution of dispersion distances is Gaussian, but still the expansion dynamics of satellite-forming biofilms is indeed faster than linear (figure 4B). On the other hand, it is also significantly slower than exponential growth (Supplementary figure S1). So, satellites form and then merge with the main colony and, thereby, limit the overall growth of the population. Nevertheless, our model suggests that in long term, the system will keep growing super-linearly, as the number of jumps (i.e. initialization of satellites) scales with the biofilm surface. In other words, as the growth of the main colony is linear and the distribution of migration distances is unchanged over time, new satellites will keep forming and the main colony will never catch up with all the satellites. However, it should be noted that our model does not encompass spreading strategies on very long time scales, as the model ignores the effects of, e.g., nutrient depletion and time-varying jumping distributions.

In our experiment, some of the main colonies of mutants produced very rough surfaces. Whilst the flagella knockout mutant’s surface (Δ*fliC*) is overall smooth, the other mutants have several small protrusions of their surfaces (figure 2B). There are a few factors that can possibly contribute to the roughness of the surfaces. The first is that even for the mutants, which are less motile than wt (figure 1C+D), individual surface cells escape occasionally. However, they travel significantly shorter distances than wt cells do. If this is the case, the satellites will quickly be re-absorbed into the main colony. In other words, the mountainous surface could be explained by motility with short dispersals (small *σ*). This is also in accordance with the fact that higher agarose concentrations (*>* 0.3%) result in smoother surfaces (figure 3). Another possibility is the nutrient driven instability of the surface growth. Several studies have shown how surface roughening instabilities are reinforced by the resulting local nutrient depletion (e.g.^48^). The nutrient-driven instability also explains the (pseudo-fractal) broccoli-like morphology recently reported for 3D *E. coli* biofilms^49^. It is possible that such instability are relevant, especially in the later stage of biofilm development. This effect is not included in our model, since it does not consider space limitations and nutrient depletion separately.

Whilst the model compares well with our experimental results with respect to the distributions of satellites, main colony volumes, and distances to satellites (figure 1C-E vs. Supplementary figure S4B-E), it fails to reproduce the distributions of the number of satellites (figure 1B vs. Supplementary figure S4A). We conclude that with only two parameters (*σ* and *k*_*s*_), we cannot cover the varieties of the natural system, i.e., the rare successful escapes. We speculate that this could be related to the variance in flagella abundance, even among cells with the same genotype. It is well-known that the expression of flagella is highly heterogeneous, i.e., there is large stochastic variations in number of flagella. This has been suggested to be an evolutionary favorable strategy, as the expression of flagella is very energy consuming^50^, such that motility is retained on colony level even when motility is limited on a cellular level.

Except for the mutant that did not express flagella, the studied strains lacked structures related to adhesion and aggregation processes rather than to motility. In general, dominant aggregation phenotypes are expected to hinder motility. Hence, it is reasonable to expect that when these processes are suppressed, the motility will increase. However, in our experiments, all the mutants presented reduced motility compared to their parental strain. This demonstrates the complexity of the bacterial motility phenotype. Indeed, vast amount of literature indicate the complex interdependence of motility and surface structure genes. Over-expression of the *handshake* protein antigen 43 (*flu*) has been reported to impair motility, not because of increased aggregation but due to interfering with the expression of flagella^51^. Similarly, the constitutive expression of type I fimbriae (*fim*) compromises the motility of bacteria by reducing the expression of flagellin^52^. The expression of curli (*csgAB*) and colanic acid (*cps*) are promoted by regulatory networks that down-regulate motility^53–55^, therefore they are normally expressed in a complementary manner. It should be noted that similar motility has been reported between wt and Δ *f lu*^51^ and between strains with and without type I fimbrial expression^56^, while this was not the case in our hands (figure 2).

Taking our results together, we suggest that satellite-formation allows bacteria to colonize unknown horizons faster (super-linear over time), whilst still retaining the tight bindings of the biofilm. Such spreading behavior can be advantageous, when invading complex environments, such as competing microbial communities, soils, or mammalian tissues. While we observed the phenomenon in a rather narrow range of the agarose concentrations, in nature it is likely that there are large gradients in the restriction of motility by the complexity of the matrix. If so, spreading through a semi-soft matrix using a combination of growth and occasional migrations, is a strategy, which ensures both fast invasion and stable occupation.

While the flagella mutant’s loss of satellites indicates the importance of flagella, more research is needed to explore exactly how rare migratory events are enabled. For example, is it the inhomogeneity of gels that sometimes allows a cell to swim over long distances before being trapped? Or is it the cells that happen to express the relevant motility genes in high enough copy numbers to overcome the gel’s resistance? Revealing the origin of the stochasticity behind rare excursions, may highlight some unknown mechanisms of robust invasion of bacteria in complex environments.

## Materials and methods

### Bacterial strains and culture media

The *Escherichia coli* strains used in this study are derivatives of the wild-type MG1655 (K-12 F-lambda-ilvG-rfb-50 rph-1)^57^ and are all listed in Table 1 and detailed in Ref.^58^ and references therein. However, the non-flagellated Δ *f liC*, (Δ *f liC*::kan) mutant (NM109), was constructed specifically for this study through P1 transduction by moving Δ *f liC*::kan from JW1908-1 (Keio collection^59^) to MS613 followed by selection on kanamycin^60^. All strains were transformed with the plasmid pVS132 carrying an Isopropyl *β* -D-1-thiogalactopyranoside (IPTG) induced yellow fluorescent protein (YFP) and ampicillin resistance for selection^61^ following protocol the of^62^.

Experiments were done using either of the two growth media: The rich Luria-Bertani (LB) composed of 1% tryptone, 0.5 % NaCl and 0.5 % yeast extract or M63 minimal media consisting of 20% 5*×*M63 salt^65^, 1 *μ*g/ml B1, 2·10^−3^ M MgSO_4_ and supplemented with 20 *μ*g/ml glucose (M63+glu). Both media were supplemented with 100 *μ*g/ml ampicillin, unless otherwise stated.

### Growth rate measurements

*E. coli* strains were grown overnight at 37°C while shaking in either LB or M63+glu medium, before diluting 1,000× in fresh medium without antibiotics. Initial OD_600_ value (NanoPhotometer^®^ C40) was measured before incubation (37°C) and repeated every hour.

### Swimming assay

Prior to experimentation, 96 mm dishes with 25 ml of either LB or M63+glu and various agarose concentrations: 0.2%, 0.3% or 0.4% (w/v) were prepared and left to dry for 18 hours on the bench. In parallel, strains were grown overnight while shaking (37°C) in LB before inoculating 1 *μ*l in the middle of each dish. After 24 hours of incubation (37°C), the range diameters were measured.

### 3D bacterial biofilms

*E. coli* strains were grown overnight at 37°C while shaking in LB medium (OD_600_ ≈ 4), before performing three consecutive thousand-fold dilutions until reaching a concentration of ≈10^3^ cells/ml. Then cells were embedded in a transparent semisolid matrix using the following procedure: First, media with 0.2%-0.6% (w/v) agarose (SERVA cat. no. 11404) was melted (in a microwave) and shaken rigorously to ensure homogeneity. To minimize evaporation, we opened new 25 ml bottles of media and agarose each time and made sure to avoid boiling. Secondly, media with agarose was aliquoted into 1 ml tubes and placed in a block-heater at 55°C. After approximately 20 min, when the mixture had reached 55°C, it was supplemented with ampicillin (100 *μ*g/ml) and 0.5 mM IPTG (for YFP induction). Then, 10 *μ*l of the diluted overnight culture were added and the mixture was immediately – to prevent untimely gelification and heat shocks – poured into a petri dish (WillCo HBST-5040) with glass bottom. This results in a final concentration of ≈10 cells pr. well (i.e 10 cells/ml). After a few minutes the mixture had solidified and the well was incubated upside-down (37°C) for 13 hours (LB) or 15 hours (M63+glu) for all agarose concentrations: 0.2% to 0.6% (w/v). The different incubation times were chosen to balance different media-dependent growth rates (Supplementary figure S1). Using these small volumes, the matrix thickness was less than 400 *μ*m, which ensures a minimum of oxygen depletion. To check this further, we compared the number of satellites pr. biofilm with the position in the well (z-direction) and found no obvious correlation. However, the cost of this thin layer is a high risk of the colonies growing onto (and spreading fast over) the surface. Therefore, we discarded all wells, with colonies where this kind of growth had happened.

### Imaging of 3D bacterial biofilms

3D biofilms in microwell dishes were imaged with a laser-scanning confocal microscope (LSCM) (Leica, SP5) and a 20*×* air immersion objective (Nplan,L20*×*,0.40corr∞). YFP was excited by an argon laser with a wavelength of 514 nm and emission was collected around 550*±*30 nm.

#### Morphology measurements

For snapshots of 3D biofilms, we collected z-stacks that captured the entire colony with (x,y)-resolution of 1.52 *μ*m/pixels and optimized z-resolution of 1.33 *μ*m. The total imaging time of the biofilm was on the order of few minutes.

### Image processing

As the typical penetration depth of a LSCM is around 100 *μ*m^66^ and the dense colonies were of the order of 200 *μ*m, the part of the biofilm furthest away in the scanning-laser direction (i.e. z-direction) suffers from strong distortions. Therefore, the following analysis was restricted to the half-colony closest to the LSCM objective by cropping the collected fluorescence z-stack using a custom-made Fiji^67^ routine. 3D image segmentation was done with BiofilmQ^68^ using the Otsu method for thresholding. Lastly, background noise was removed by eliminating outlier voxels of clusters smaller than 11 *μ*m^3^. The following BiofilmQ parameters were exported for subsequent analysis: Convexity, number of satellites, volumes, distances between center-of-masses and nearest-neighbour objects.

The *in silico* biofilms was processed in the same manner, but skipping the thresholding step (as they are binary *per se*).

### Modified Eden growth model

To reproduce the features of the obtained experimental results a modified Eden growth model was implemented. In a three-dimensional cubic lattice of linear system size *L*, each lattice site can be occupied by at most one bacterial cell. Each cell can grow at a rate *k* if there is at least one empty site among their 6 next nearest-neighbour sites. On top of this, if 3 or more of the next nearest sites are empty, then in addition to the growth, the cell can jump to another location. This is implemented as a jump to a new randomly chosen location that happens at a rate *k*_*s*_. This is implemented by counting the number of the cells that can only grow *N*_1_ and the number of cells that can both grow and swim *N*_2_ at each update and by applying a Gillespie algorithm.

The initial condition is a single cell placed in the center of the system and then the following steps are iterated:

1. Find all the cells in the lattice and count the number of their empty nearest-neighbours. Among them, a cell that has 1-3 available neighbouring sites belongs to the population *N*_1_, and otherwise, the cell belongs to the population *N*_2_.
2. Define an array containing all empty neighbouring sites, corresponding to the surface of the colony.
3. Compute the total event rate *T* = *N*_1_ · *k* + *N*_2_ · (*k* + *k*_*s*_), and determine the duration to the next event as *τ* = − ln(*r*)*/T*, where *r* is a random number from a uniform distribution (r = *U* ⊆ (0, 1)). Proceed the time by *τ*.
4. Draw a random uniform number, *a*, from a uniform distribution between zero and one (a= *U* ⊆ (0, 1)) and determine which event to happen by the following procedure:
  4.1. If *a >* (*N*_1_ + *N*_2_) · *k/T* a growing event happens. Choose a random surface site and add a new cell to it.
  4.2. Otherwise, a swimming event happens. Choose a random cell of the sub-population *N*_2_. Generate three random Gaussian distributed numbers of zero mean and standard deviation *σ*, to find the new position of the cell. If the new cell position is empty, update the cell position. If the new position is already occupied by another cell, draw a new position from the above-mentioned procedure until finding a new empty position.
5. Return to (1).

Pseudo-code is provided in the Supplementary text.

### Statistics

All mean values are given as (mean *±* SD) unless stated otherwise and only when data are tested against the null hypothesis that it is normally distributed.

## Data availability

The code to generate the simulated data is available in Zenodo with DOI: 10.5281/zenodo.7414919.

