## Supplementary figures for "Motility mediates satellite formation in confined biofilms"

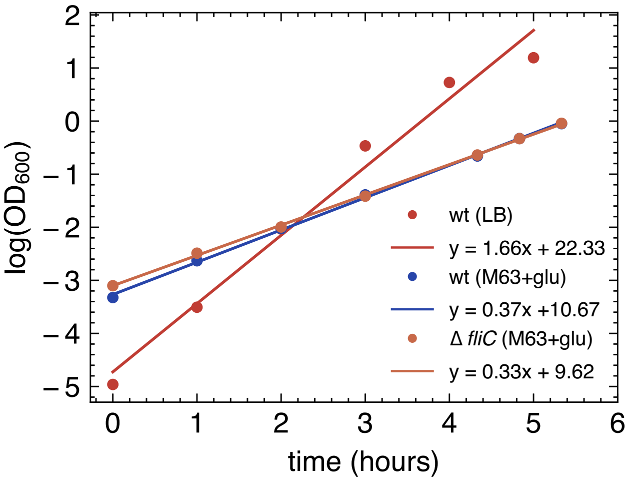


**Figure S1:** Growth curves for parental strain (wt) and flagella mutant (Δ*fliC*). Plot contains the experimental data (circles) together with the exponential fit (full lines). The doubling time is found as the $ln(2)$ divided by the slope of the growth curve (full lines). The resulting doubling times are 27.8 min (wt, LB), 68.4 min (wt, M63+glu), and 72.8 min (Δ*fliC*, M63+glu).

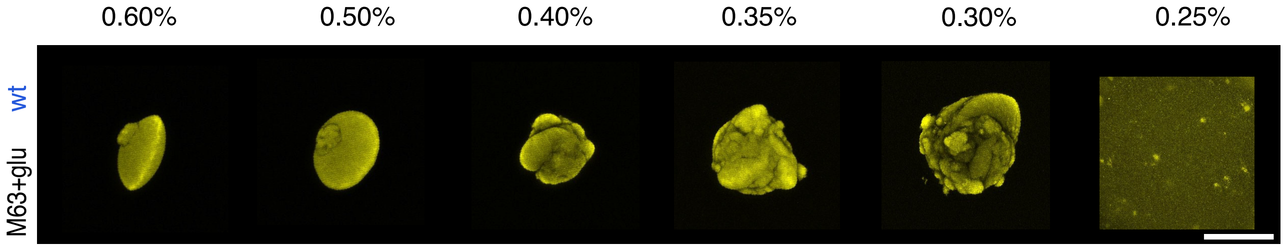


**Figure S2:** Effect of matrix elasticity on 3D biofilm morphology. Examples of pseudo-colored fluorescent wt biofilms (maximum intensity projections) grown in minimal medium (M63+glu) at various agarose concentrations. The scale bar corresponds to 200 $\mu$m. At low agarose concentration (0.25%) cells swims through the media and at high concentrations ($> 0.50\%$) biofilms' have an oblate shape and a smooth surface.

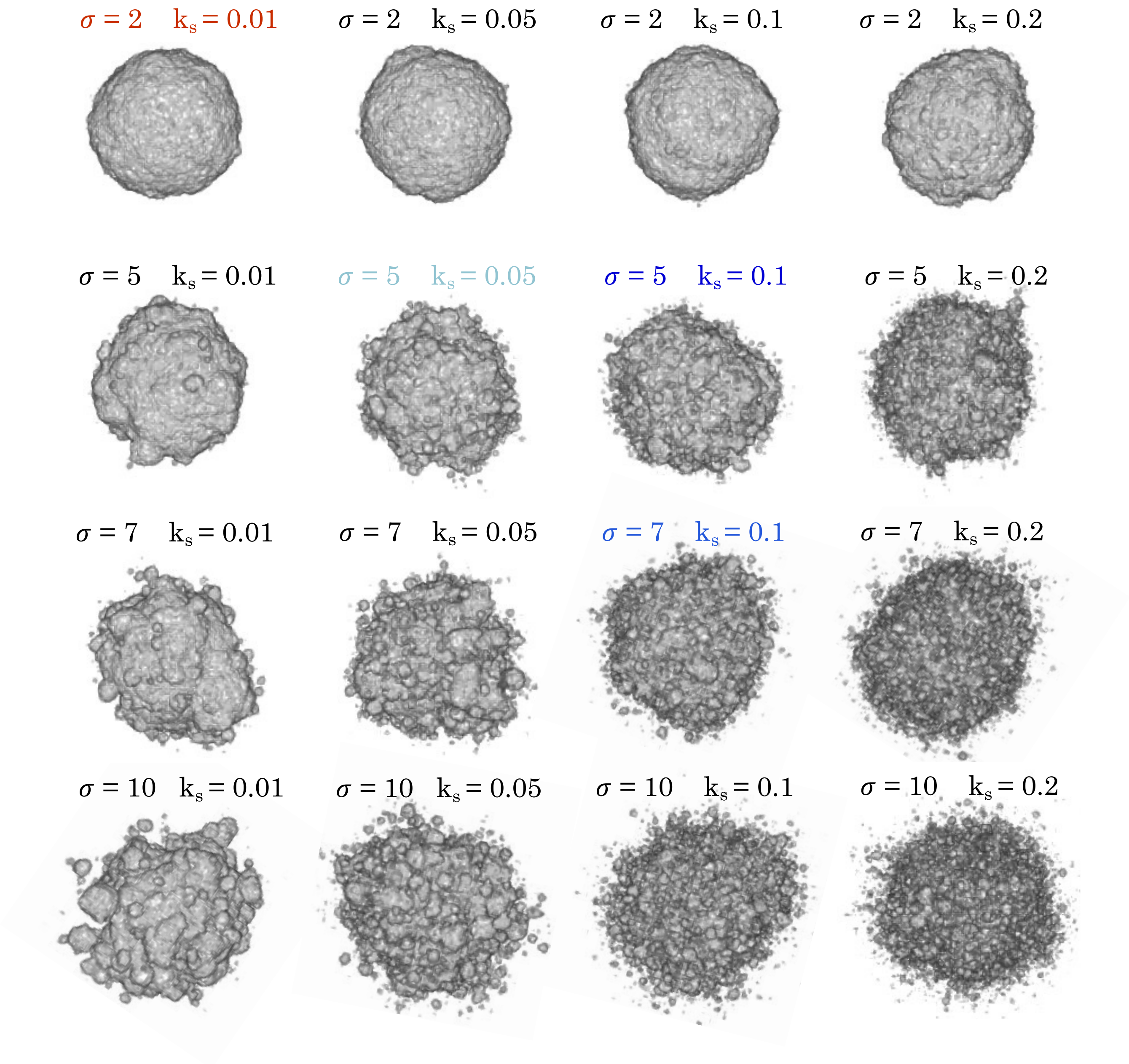


**Figure S3:** Effect of frequency, $k_s$, and distance, $\sigma$, of jumps in the simulations of 3D biofilms after $10^6$ division/jump events. The color-coding is the same as in figure 4A in the main text.

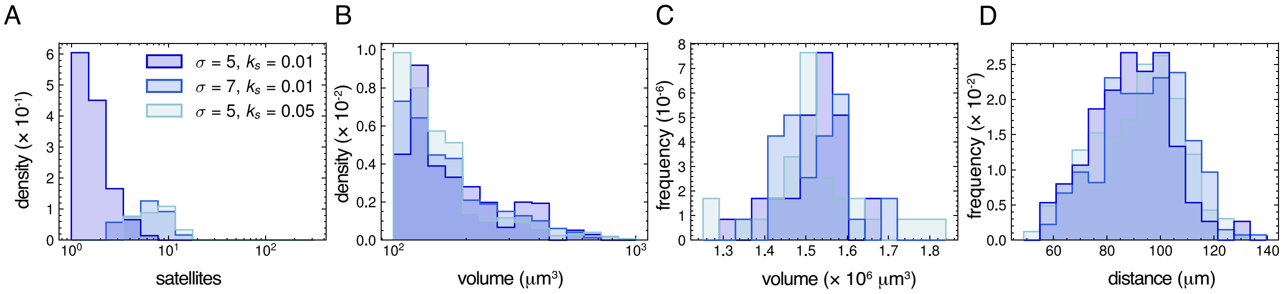


**Figure S4:** Satellite morphology in 3D biofilms. Simulated 3D biofilms (N=30) for 3 different sets of $\sigma$ and $k_s$ (N=30). Each lattice site corresponds to 1 $\mu$m and the color-coding is the same as in figure 4A in the main text. **B:** Distribution of of the number of satellites pr. biofilm (log-scale). **C:** Distributions of satellite volumes (log-scale) in either $\sigma$ = 5 and $k_s$ = 0.01 (N=61), $\sigma$= 7 and $k_s$ = 0.01 (N=223), and for $\sigma$= 5 and $k_s$ = 0.05 (N=271). **D:** Distribution of volumes of the main colonies. **E:** Distributions of distances from the center-of-mass of satellites to the center-of-mass of the main colony.

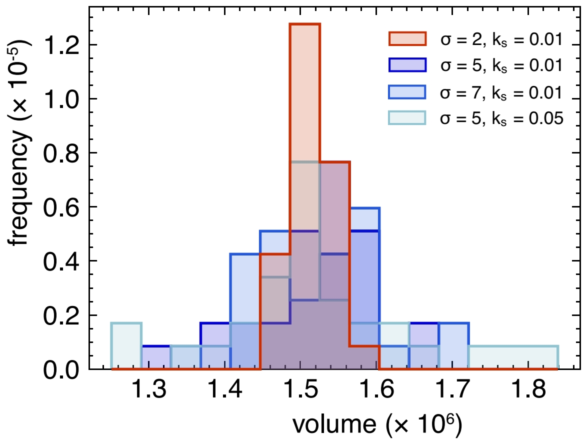


**Figure S5:** Total 3D biofilm volumes for simulated 3D colonies (N=30), where each lattice site corresponds to 1 $\mu$m. Color-coding is the same as in figure 4A in the main text.
