## Supplementary code for "Motility mediates satellite formation in confined biofilms"

### Computational Tools

**Input:** Lattice size $L$, growth rate $k$, swimming rate $k_s$, standard deviation $std$ and an average $\mu$=0 of a Gaussian distribution and the total number of events $P$

**Result:** Aggregate of cells in a 3D lattice

1 Generate a 3D lattice of dimension $L$ with empty sites $Lat(i, j, k) = 0$ and occupied sites $Lat(i, j, k) = 1$;

2 Place a single cell in the center of the lattice

$$Lat\ (L/2, L/2, L/2) = 1$$

**for** $p \leftarrow 0$ **to** $P$ **do**

3 Define an empty colony surface, $S$;

4 Define an empty swimmers array, $swim$;

5 Set the number of each cell types, $N1$, $N2$ to zero;

6 **foreach** *occupied site* $Lat(i, j, k) = 1$ **do**

7 Find and count the number of empty near neighbours $\beta$

$$Lat(i \pm 1, j, k) = 0 \quad Lat(i, j \pm 1, k) = 0 \quad Lat(i, j, k \pm 1) = 0$$

Add the available sites position to the surface of the colony $S$.

8 **if** $\beta > 3$ **then**

9 Add an element to $N2$ ;

10 Save the cell position in $swim$;

11 **else if** $0 < \beta \leq 3$ **then**

12 Add an element to $N1$ ;

13 **end**

14 Find the total event rate $T = N1 \cdot k + N2 \cdot (k + k_s)$;

15 Determine which event happens at time $t = -ln(r)/T$, with $r = U \subseteq (0, 1)$, by trowing a random number from an uniform distribution $a = U \subseteq (0, 1)$ ;

16 **if** $(N1 + N2) \cdot k/T < a$ **then**

17 Chose a random surface site $S(m, n, l)$ and update the colony by setting a cell in that position $Lat(m, n, l) = 1$;

18 **else**

19 Throw three Gaussian numbers to determine the final position of the cell: $x_g = P(x) \quad y_g = P(y) \quad z_g = P(z)$;

20 Select an element of the swimmers array $Lat(m', n', l')$;

21 **if** $Lat(m' + x_g, n' + y_g, l' + z_g) = 0$ **then**

22 $Lat(m', n', l') = 0$;

23 $Lat(m' + x_g, n' + y_g, l' + z_g) = 1$;

24 **else**

25 Go to 19;

26 **end**

27 **end**

28 **end**

**Algorithm 1:** 3D Eden Growth Model with swimmers
